# Effects of dietary replacement of broiler litter with *Mucuna pruriens utilis* forage and seed meal on performance, carcass characteristics, blood biochemical and physiological parameters in indigenous goats

**DOI:** 10.1101/421313

**Authors:** Doctor M.N. Mthiyane, Nozipho P. Gamedze, Abednego M. Dlamini, Arno Hugo, Ignatius V. Nsahlai

## Abstract

The productivity of indigenous goats in Africa is constrained by shortage of protein-rich feed especially in winter. This study investigated the nutritional value of mucuna forage (MF) and seed meal (MSM) as alternative protein sources for indigenous goats. Mucuna was planted in 3 parallel and adjacent fields and its foliage nutritional composition determined at 4, 8, 12 and 16 weeks after planting (WAP). MF was harvested at 14 WAP whilst mucuna pods were harvested at 28 – 30 WAP, shelled and the MSM chemically analysed. In a completely randomised design (CRD), 20 indigenous goats were randomly offered 5 treatment diets with, respectively, 0%, 25%, 50% and 100% MF and 100% MSM replacing broiler litter (BL), each with 4 replicates, for 82 days. Both mucuna foliage DM and CF contents increased (P < 0.001) whilst foliage CP, EE and ash contents decreased (P<0.001) with maturity. On the other hand, MSM contained high DM (90.7%), EE (3.7% DM) and CP (26.0% DM) but low CF (9.7% DM) and ash (5.5% DM) contents. Both body weight gain (BWG) and feed conversion efficiency (FCE) were not influenced by dietary mucuna incorporation (P > 0.05). However, dietary MF, particularly at the 100% level, decreased goat feed intake (FI) (P < 0.001) whilst 100% MSM increased (P < 0.001) this parameter. Mucuna had no effect on all carcass characteristics (P > 0.05) but increased (P < 0.05), particularly at the 100% MSM level, hot carcass weight and dressing percentage. There were no effects of mucuna on all biochemical and haematological indices (P > 0.05), except for the increase in serum glucose (P < 0.05). In conclusion, the optimal stage for harvesting and utilisation of MF is between 12 and 16 WAP and both MF and MSM, particularly the latter, are rich alternative protein sources for indigenous goats.

## Introduction

The world goat population is ∼921 million, >90% of which are found in developing countries [1]. Of these, ∼352 million goats are in Africa [2], >90% of which are owned by smallholder farmers [3]. The major concentration of goats in Africa is in the sub-Saharan region with 60% of the total goat population representing ∼80 indigenous breeds raised under various production systems [3]. Indigenous goats (*Capra hircus*) play important roles mainly in terms of food and nutrition security, poverty alleviation, and cultural life of smallholder rural-based farmers and societies in Africa [4, 5, 6]. They provide immediate benefits in the form of meat, milk, leather, cash income, manure and other miscellaneous products to the poor in communal areas [5, 7] and also play a pivotal role in traditional ceremonies [6, 8]. They generally have small to medium frame with short horns and a variety of coat colours [4, 9]. Unlike the Boer and dairy goats, indigenous goats remain rather unimproved local varieties [10] and, unlike cattle, are relatively neglected in many African societies, despite their myriad advantages. These varieties are not defined but usually associated with the geographical area in which they are found and are mostly valued for their hardiness, resistance to endemic diseases, early attainment of maturity, and possession of a unique ability to adapt to a variety of climatic conditions and for reproducing under low input systems [3, 9]. They can also efficiently survive on available shrubs and trees in arid and semi-arid environments where hardly any crops can be successfully grown [11]. They also provide higher off-take than cattle because of their shorter generation interval and higher prolificacy. Annual off-take can be as high as 60 % [12] and, in favourable environments, they can attain twinning rate of >70% by second kidding [13, 14].

Globally, whilst goat meat consumption is less than beef [15], the goat industry has become an important livestock enterprise due to a growing demand for chevon [16, 17]. The rise in chevon demand is ascribed to its general consumer perception as being leaner and having less cholesterol content, greater polyunsaturated (PUFA) to saturated (SFA) fatty acid ratio and favourable sensory characteristics in comparison to beef, lamb/mutton and pork [17, 18]. SFAs, as opposed to their PUFA counterparts, are generally associated with the worldwide proliferation of modern diseases of diabetes, cardiovascular disease, cancer and others [19, 20, 21]. The rise in chevon demand, therefore, presents an opportunity for rural smallholder farmers in Africa to contribute in the fight against human diseases of global significance and, in the process, reap benefits in terms of economic development by connecting with goat value chains. For example, with a predominantly rural (80 %) human population of 1.13 million that grows at approximately 1.2 % annually [22], about 70% of which is dependent on subsistence agriculture for their livelihoods [23, 24, 25], and with 73 % of the population living below the national poverty line [22], the Kingdom of Eswatini (Swaziland) can enormously benefit from improvements in goat production. Currently, the goat population in this Southern African country is approximately half a million [26] and rearing of the animals is practiced at subsistence level by a large section of the rural population of smallholder farmers [3].

The productivity of indigenous goats in Southern Africa is, notwithstanding, constrained by a shortage of good quality feed especially in winter. In this regard, indigenous goat nutrition in this region of the African continent is largely based on communal natural pastures and rangelands in which available tropical forages are highly fibrous and low in protein during winter season. Consequently, this compromises goat productivity [27] and predisposes the animals to diseases and parasites and ultimately dearth in extreme cases [28, 29]. Also, abundant agro-industrial by-products [e.g. sugarcane tops, of which The Kingdom of Eswatini produced ∼920 000 metric tonnes in 2016/2017 (20% of sugarcane production, on wet basis) [30]] are deficient in and require supplementation with proteins and minerals [31]. Unfortunately, commercially available dietary supplements based on conventional protein sources such as soya bean meal are too expensive and unaffordable by most smallholder farmers. Hence the urgent need for a search for cheaper locally available yet nutritionally adequate alternative feed supplements.

Broiler litter (BL) is one of the most extensively used and intensively investigated cheap alternative protein (15% – 35% CP) and mineral rich supplementary feedstuffs for goats and other livestock in Southern Africa [31, 32, 33]. The utilisation of BL is, however, limited by the presence of human pathogens [34], pesticides and drug residues [35] and heavy metals [36] which might pose health hazards to livestock and human consumers of livestock products. One safer and sustainable strategy of improving goat productivity involves supplementation with protein-rich tropical leguminous forages such as mucuna.

Mucuna or velvet bean [*M. pruriens* (L.) DC var. *utilis*; *Fabaceae* family] is a vigorous annual climbing tropical legume indigenous to Southern Africa [37]. It is cultivated in Africa, America, Asia and the Pacific Islands [38], is adapted to tropical environments with tolerance to higher environmental temperatures and other abiotic stresses [39], exhibits aggressive growth and is not suppressed by weeds [40]. Its vines and foliage can be used as pasture, hay or silage for ruminants while pods and seeds can be ground into a meal and fed to both ruminants and non-ruminants [41]. Mucuna seeds are rich in proteins (23 – 35%) [42], with high digestibility (70 – 80%) [43] and an essential amino acid composition comparable to the FAO/WHO requirement [44]. They are also rich in minerals and essential fatty acids [45, 46] and are therefore regarded as a good source of food for humans [38], ruminants [47] and non-ruminants [48, 49, 50]. Otherwise, mucuna seeds are well-known for their high content of 3,4-dihydroxy-L-phenylalanine (L-DOPA), the most effective symptomatic medication of Parkinson’s disease and the gold standard in the treatment of the disease [51, 52].

The presence of L-DOPA, the concentration of which is highest in the seeds [53], has been the main hindrance in the widespread utilization of mucuna as animal feed. However, since goats have the ability to detoxify secondary compounds which enables them to consume plants or plant parts with a high content of chemical defenses [54, 55], it is unlikely that L-DOPA would cause toxicity problems to this ruminant livestock species. Whilst L-DOPA has been reported to be extensively metabolised by rumen microorganisms [56, 57], there are limited studies that have previously investigated the nutritional value of the L-DOPA-containing mucuna for indigenous goats [47, 58, 59, 60]. It was thus hypothesized that dietary incorporation of MF and MSM would improve performance and carcass characteristics without detrimentally affecting the serum biochemical and haematological indices in the goats. The objectives of the study were therefore to determine (a) the optimum stage of harvesting of MF in terms of its nutritional value, and (b) the effects of dietary MF and MSM incorporation on performance, carcass characteristics, serum biochemical and haematological indices in indigenous goats.

## Materials and methods

### Location of study, source and preparation of materials

Mucuna planting and the feeding trial were conducted at the University of Eswatini (UNESWA) Luyengo campus farm located in the upper Middleveld of The Kingdom of Eswatini (coordinates: 26° 32’ south and 31° 14’ east, altitude: 600 – 800 m above sea level; mean maximum and mean minimum temperatures: 23 °C and 11 °C, respectively). The annual rainfall ranges from 850 mm – 1000 mm [61], most of which occurs between October and April. Chemical analyses of experimental diets were performed in the Nutrition Laboratory of the Department of Animal Science, Luyengo Campus. Analyses of haematological and biochemical indices were conducted at Lancet Laboratories in Mbabane.

Mucuna seeds were obtained from the first author as well as from Malkerns Research Station in The Kingdom of Eswatini. They were scarified by immersion in boiling water for 20 seconds to facilitate germination. Land was prepared using a tractor. A total of 7 × 50 kg bags of NPK 2:3:2 (37) fertiliser was applied to the soil. Mucuna beans were planted in October 2016 immediately after the first rains at a spacing of 22 cm intra-row and 90 cm inter-row in 3 parallel and adjacent fields using a tractor mounted planter. During the first 3 WAP, the fields were irrigated using sprinklers in order to ensure seedling germination and establishment after which they were rain fed. Weeding was done by hand 3 weeks after germination. It was repeated as and when necessary depending on weed growth. Seedling growth performance was regularly observed and seeds that did not germinate were manually replaced through gap filling.

MF was harvested by hand at 14 WAP and dried under shade for 2 weeks with daily manual turning using a hand-held fork to facilitate drying. To determine the effect of stage of growth on the nutritive value, mucuna foliage (leaves only) was sampled at 4, 8, 12 and 16 WAP and then taken to the Nutrition Laboratory for drying (65 °C), milling (1 mm) and chemical analysis. On the other hand, dry mucuna pods were harvested at 28 – 30 WAP and kept on a shed floor at room temperature until they were completely dry to allow manual shelling. Samples of dry mucuna seeds were taken and milled (1 mm) for chemical analysis.

Sugarcane tops were obtained from Dalcrue Farm and dried for 2 weeks under sunlight with daily manual turning using a hand-held fork to facilitate drying. Hominy chop, mineral-vitamin premix and salt were supplied by Feedmaster (Pty) Ltd (Eswatini). Molasses was obtained from Royal Swaziland Sugar Corporation (RSSC) at Mhlume whilst BL was from Matsapha Correctional Centre (Livestock Section). MF, mucuna seeds, sugarcane tops and BL were then milled (13mm sieve) using a tractor-mounted miller. Twenty intact male indigenous goats (12 – 18 months old; average initial liveweight = 19.6 ± 2.7 kg) were sourced from a local farmer at Malindza area in the Lubombo region of Eswatini. They were dewormed using Dectomax (1ml/10kg BW) and vaccinated with Berenil and Terramycin LA. Further, they were vaccinated against pulpy kidney disease. Dipping was done once before the trial using Paracide Dip to prevent tick-borne diseases. Further treatments were applied when necessary.

### Experimental design, animals and diets

Following a 14-day adaptation period, 20 goats were used in an 82 day-long CRD experiment involving 5 dietary treatments with 4 replicates each. The individual goat was the experimental unit. The animals were randomly assigned to experimental diets based on body weight. Iso-energetic and iso-nitrogenous sugarcane tops and hominy chop-based diets, in meal form, formulated to meet the nutritional requirements of goats as recommended by the NRC [62] and to which graded levels of MF [0% (Control), 25%, 50% and 100 %] and MSM [100%] were sequentially added in replacement of BL (Table 1), were used. The experimental diets were mixed at Arrowfeeds (Pty) Ltd (Eswatini).

**Table 1.** Ingredient and nutrient composition (%) of experimental diets.

| Ingredient | Mucuna level |  |  |  |  |
| --- | --- | --- | --- | --- | --- |
|  | Forage |  |  |  | Seed meal |
|  | 0% | 25% | 50% | 100% | 100% |
| Sugarcane<br>tops | 33.0 | 33.0 | 33.0 | 33.0 | 33.0 |
| Hominy<br>chop | 20 | 20 | 20 | 20 | 20 |
| BL | 36.0 | 27.0 | 9.0 | 0.0 | 0.0 |
| Mucuna | 0.0 | 9.0 | 27.0 | 36.0 | 36.0 |
| Molasses | 10.0 | 10.0 | 10.0 | 10.0 | 10.0 |
| Salt | 0.5 | 0.5 | 0.5 | 0.5 | 0.5 |
| Mineral-vitamin premix | 0.5 | 0.5 | 0.5 | 0.5 | 0.5 |
| *Analysed chemical composition (%)* |  |  |  |  |  |
| DM | 90.0 | 89.0 | 89.9 | 90.1 | 89.8 |
| CP | 15.5 | 15.8 | 16.0 | 15.7 | 16.4 |
| CF | 19.9 | 23.4 | 22.4 | 22.5 | 13.47 |
| EE | 0.6 | 2.2 | 2.8 | 2.9 | 3.6 |
| Ash | 6.9 | 5.0 | 4.8 | 4.0 | 4.0 |

<sup>a</sup>Mineral-vitamin premix supplied the following per kg of complete feed: vitamin A (retinol acetate) 4000 000 IU, copper 10g, manganese 30g, zinc 70g, iodine 0.8g, cobalt 0.3g and selenium 300 00mg.

### General management

Goats were kept in wooden pens (1 m × 2 m; 1 goat/pen) built inside a well-ventilated brick walled house with sides open to natural light, concrete floor and a roof of corrugated iron sheets. Before commencement of the experiment, pens were thoroughly cleaned and disinfected with an antiseptic (Jeyes fluid and Vet Range). Sawdust was used as bedding and was regularly changed at 2-week intervals. Goats were allowed *ad libitum* access to feed and water and were inspected daily for any health-related problems. Their diets were offered as a total mixed ration twice daily at 08h00 hrs and 17h00 hrs.

In the evening of the last day (day 82) of the experiment, all goats were deprived of feed for 12 hrs during which they were provided with clean water only. They were humanely slaughtered in the morning of day 83 at the UNESWA Farm abattoir following MOA [63] procedures. After slaughter, bleeding and removal of hides, evisceration was done in preparation for the measurement of hot carcass and cold carcass parameters.

### Measurements

Goats were individually weighed at the beginning (day 0) and then fortnightly until the end of the experiment. Also, feed offered and left was weighed per pen daily in order to calculate average FI. FI was therefore measured as feed offered minus refusals. FCE was calculated as BWG/FI.

The hot carcass of each goat was weighed after removal of the head, hooves and viscera. Carcasses were then kept at room temperature for 3 hrs to ensure blood drainage before they were transported into a cold room (2 °C). After 24hrs of chilling, cold dressing percentage (%) and chilling loss were calculated as follows:

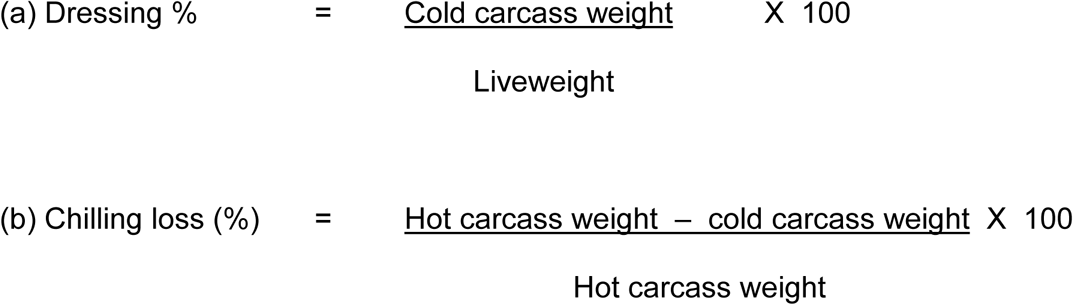

Blood samples (10 ml) were collected (into vacutainers containing EDTA, for haematological parameters, and vacutainers without anticoagulant, for biochemical analysis) on day 82 of the experiment by bleeding the goats through the jugular vein. Vacutainers were immediately placed on ice and then taken to Lancet Laboratory for haematological analysis.

All animals were cared for according to the animal care and welfare guidelines of the MOA [63] and UNESWA Department of Animal Science Board that approved the protocol used in the experiment.

### Chemical assays

The DM (930.15), ash (942.05), CP (954.01), EE (920.39) and CF analyses of mucuna foliage, MSM and experimental diets were performed according to AOAC [64] procedures. CP was calculated using N × 6.25. Haematological analysis was performed using the BC-5380 Mindray Auto Haematology Analyzer following the method described by Diamond Diagnostics. Serum glucose, total protein, albumin, urea and globulin were determined using the Roche Integra analyser following the Colourimetric method.

### Statistical analyses

Data on mucuna foliage nutritional composition, growth performance and carcass characteristics were analysed using the General Linear Models procedure [65]. Statistical significance was accepted based on the 0.05 level of probability. Data are presented as Least Square Means (LSM) with respective pooled SEM. Where significant differences (p < 0.05) between treatments were observed, LSM were compared using the Tukey test.

## Results

The nutritional composition of mucuna foliage and MSM is shown in Table 2. Mucuna foliage DM increased (P < 0.001) from 4 WAP (17.6%) to reach a maximum between 12 (24.2%) and 16 (23.0%) WAP. The foliage CF content also linearly increased (P<0.001) with mucuna plant maturity. In contrast, mucuna foliage CP decreased (P<0.001) from 30.6% DM at 4 WAP to around 27% DM between 12 and 16 WAP. Also, both mucuna foliage EE (P<0.001) and ash (P<0.001) contents linearly decreased with maturity.

**Table 2.** The proximate composition (% DM) of mucuna foliage at various stages of growth and seed meal at 28 to 30 WAP.

| Parameter | Mucuna |  |  |  |  | SEM<sup>1</sup> | P-value |
| --- | --- | --- | --- | --- | --- | --- | --- |
|  | Foliage |  |  |  | Seed meal |  |  |
|  | Week 4 | Week 8 | Week 12 | Week 16 | Week 28-30 |  |  |
| DM | 17.6 | 21.9 | 24.2 | 23.0 | 90.7 | 0.21 | P < 0.001 |
| CP | 30.6 | 29.6 | 26.7 | 27.4 | 26.0 | 0.22 | P < 0.001 |
| CF | 24.5 | 22.1 | 25.0 | 26.2 | 9.7 | 0.30 | P < 0.001 |
| EE | 2.5 | 1.9 | 1.4 | 0.8 | 3.7 | 0.12 | P < 0.001 |
| Ash | 9.0 | 8.5 | 6.5 | 7.7 | 5.5 | 0.10 | P < 0.001 |

<sup>1</sup>SEM based on pooled estimate of variation (n = 9).

On the other hand, MSM contained high DM (90.7%), EE (3.7% DM) and CP (26.0% DM) but low CF (9.7% DM) and ash (5.5% DM) contents (Table 2).

The performance parameters (Table 3) and carcass characteristics (Table 4) of goats fed treatment diets are presented below. Both BWG and FCE were not influenced by dietary mucuna incorporation (P > 0.05). However, dietary incorporation of MF, particularly at the 100% level, decreased goat FI (P < 0.001) whilst incorporation of 100% MSM increased (P < 0.001) this parameter (Table 3). There was no effect (P > 0.05) of mucuna incorporation on all carcass characteristics, except for the increase in hot carcass weight (P < 0.05) and dressing percentage (P < 0.05), particularly at the 100% MSM level (Table 4).

**Table 3.** Body weight gain, feed intake and feed conversion efficiency of indigenous goats fed diets supplemented with graded levels of mucuna forage and seed meal.

| Parameter | Mucuna level |  |  |  |  | SEM<sup>1</sup> | P-value |
| --- | --- | --- | --- | --- | --- | --- | --- |
|  | Forage |  |  |  | Seed meal |  |  |
|  | 0% | 25% | 50% | 100% | 100% |  |  |
| BWG (g/d) | 35.7 | 20.8 | 41.7 | 40.2 | 35.7 | 11.1 | P > 0.05 |
| FI (g/d) | 845.3 | 709.6 | 837.7 | 627.7 | 899.8 | 9.6 | P < 0.001 |
| FCR<br>(BWG/FI) | 0.05 | 0.02 | 0.03 | 0.13 | 0.01 | 0.03 | P > 0.05 |

<sup>1</sup>SEM based on pooled estimate of variation (n =4).

**Table 4.**
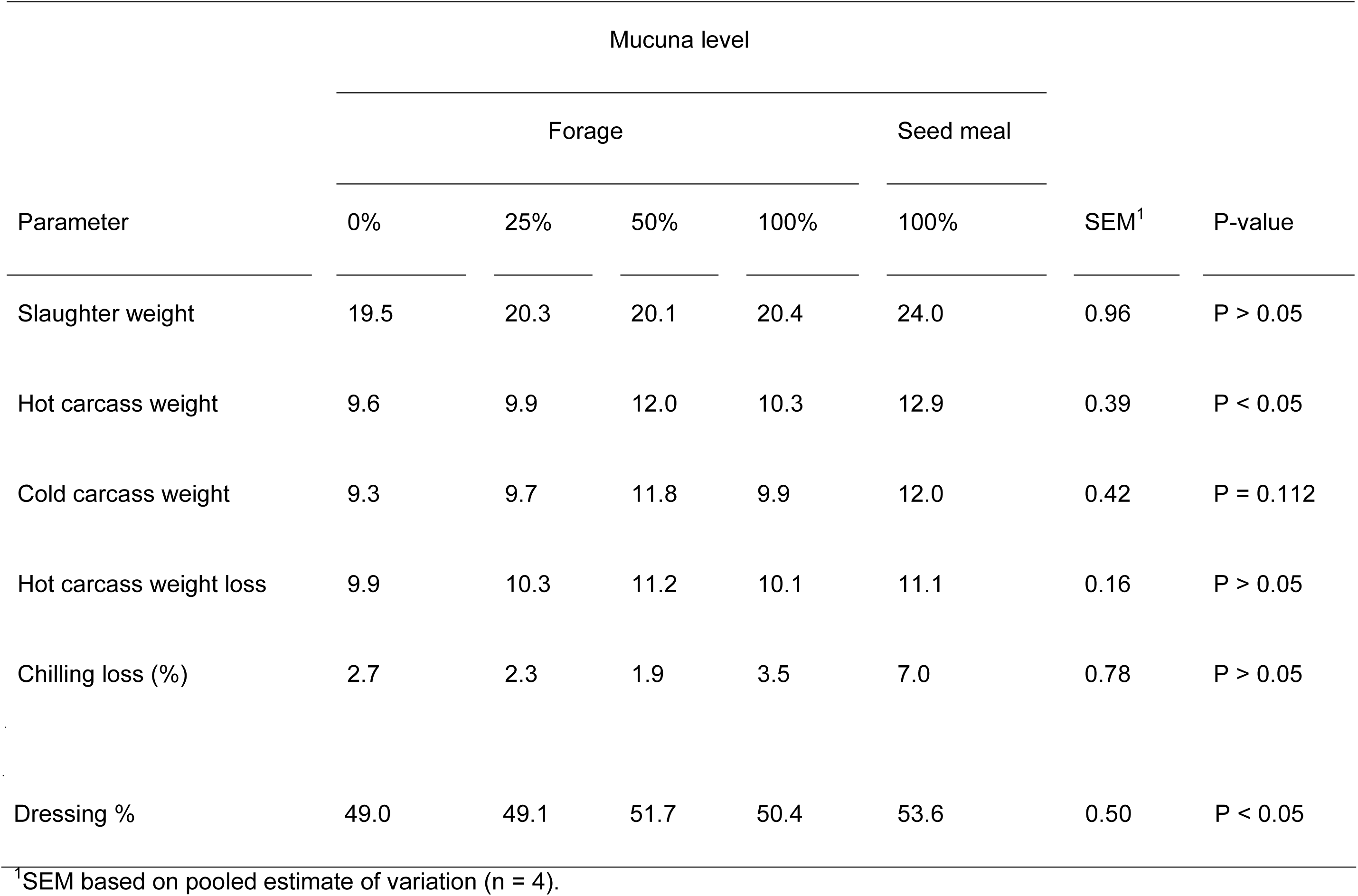
Carcass characteristics (kg) of indigenous goats fed diets supplemented with graded levels of mucuna forage and seed meal.

The influence of dietary incorporation of mucuna on biochemical and haematological indices of goats is respectively presented in Tables 5 and 6. There was no effect of dietary mucuna incorporation on all biochemical and haematological indices (P > 0.05), except for the increase in serum glucose induced by particularly the 50% and 100% MF and 100% MSM levels (P < 0.05).

**Table 5.** Serum biochemical parameters of indigenous goats fed diets supplemented with graded levels of mucuna forage and seed meal.

| Parameter | Mucuna level |  |  |  |  | SEM<sup>1</sup> | P-value |
| --- | --- | --- | --- | --- | --- | --- | --- |
|  | Forage |  |  |  | Seed meal |  |  |
|  | 0% | 25% | 50% | 100% | 100% |  |  |
| Glucose (mmol/L) | 5.7 | 4.5 | 7.1 | 7.0 | 9.0 | 0.40 | P < 0.05 |
| Total protein (g/L) | 64.0 | 60.3 | 64.7 | 63.5 | 66.0 | 2.24 | P > 0.05 |
| Albumin (g/L) | 28.8 | 24.3 | 27.0 | 26.3 | 26.0 | 1.22 | P > 0.05 |
| Urea (mmol/L) | 5.9 | 4.3 | 5.8 | 4.9 | 6.4 | 0.32 | P > 0.05 |
| Globulin (g/L) | 35.3 | 36.0 | 44.3 | 37.3 | 40.0 | 2.22 | P > 0.05 |

<sup>1</sup>SEM based on pooled estimate of variation (n =4).

**Table 6.**
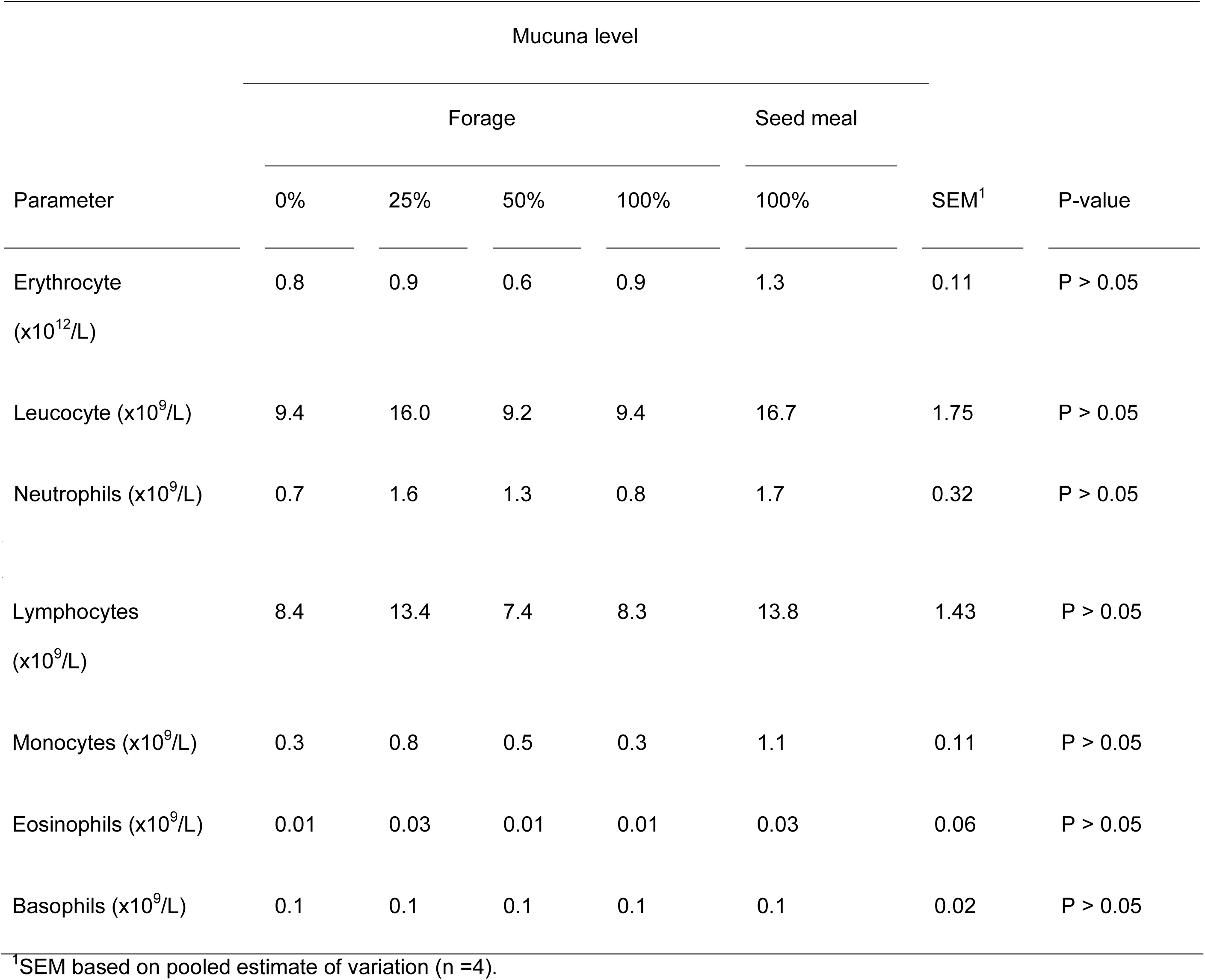
Haematological parameters of indigenous goats fed diets supplemented with graded levels of mucuna forage and seed meal.

| Parameter | Mucuna level |  |  |  |  | SEM<sup>1</sup> | P-value |
| --- | --- | --- | --- | --- | --- | --- | --- |
|  | Forage |  |  |  | Seed meal |  |  |
|  | 0% | 25% | 50% | 100% | 100% |  |  |
| Erythrocyte<br>(x10<sup>12</sup>/L) | 0.8 | 0.9 | 0.6 | 0.9 | 1.3 | 0.11 | P > 0.05 |
| Leucocyte (x10<sup>9</sup>/L) | 9.4 | 16.0 | 9.2 | 9.4 | 16.7 | 1.75 | P > 0.05 |
| Neutrophils (x10<sup>9</sup>/L) | 0.7 | 1.6 | 1.3 | 0.8 | 1.7 | 0.32 | P > 0.05 |
| Lymphocytes<br>(x10<sup>9</sup>/L) | 8.4 | 13.4 | 7.4 | 8.3 | 13.8 | 1.43 | P > 0.05 |
| Monocytes (x10<sup>9</sup>/L) | 0.3 | 0.8 | 0.5 | 0.3 | 1.1 | 0.11 | P > 0.05 |
| Eosinophils (x10<sup>9</sup>/L) | 0.01 | 0.03 | 0.01 | 0.01 | 0.03 | 0.06 | P > 0.05 |
| Basophils (x10<sup>9</sup>/L) | 0.1 | 0.1 | 0.1 | 0.1 | 0.1 | 0.02 | P > 0.05 |

<sup>1</sup>SEM based on pooled estimate of variation (n =4).

## Discussion

The nutritional composition of mucuna foliage observed in the current study was within range of values reported in the literature [66]. Also, the observed increase in mucuna foliage DM and CF contents in contrast to a decrease in CP, EE and ash contents with maturity corroborates previous findings [66, 67, 68, 69]. The mechanism underlying the age-dependent increase in foliage CF in contrast to a decrease in its CP, EE and ash contents may involve deposition of photosynthetic carbon into structural material thus diluting the metabolic pool as represented by cell contents as the plant develops [70]. Considering the contrasting CP and fibre dynamics with maturity, we determined the optimal stage for harvesting and utilization of MF under our conditions to be between 12 and 16 WAP. Hence, MF used in the current study was harvested at 14 WAP.

Regarding MSM, the observed nutritional content was also within the range of values reported in the literature for CP (20.2 – 30.3%) [44, 71, 72], EE (1.84 – 14.39%) [73], CF (1.6 – 7.9%) [74] and ash (2.9 – 4.4%) [75]. In other studies, Ezeagu *et al*. [45] found similar CP (24.5 – 29.8%) but slightly higher EE (4.7 – 7.3%) and lower CF (3.7 – 4.4%) in seeds of 12 *Mucuna* accessions. Also, Mupangwa *et al*. [76] observed rather lower CP content (16.8%) for mucuna hay compared to mucuna foliage values in the current study. The variations in nutritional content might be due to geo-climatic differences. Notwithstanding, both MF and MSM used in the current study contained rather high CP contents to qualify them for use as protein supplements in feeds for indigenous goats and other livestock.

The observed lack of effect of dietary MF and MSM incorporation on goat BWG and FCE in the current study corroborates previous studies with goats [58, 77, 78] and cows [79, 80]. Our results, however, disagree with other studies that found improved BWG [81, 82, 83] in sheep and milk yield in cows [84] fed diets supplemented with MF and MSM. They further disagree with those that found decreases in BWG [56, 60, 85], milk yield and liveweight [47] in indigenous goats and growing lambs fed diets supplemented with MSM and whole pods. The inconsistency in responses to mucuna supplementation might be related to variations in the dietary level of L-DOPA. In this connection, it would appear that almost all studies that reported negative effects of mucuna involved the seeds and/or pods rather than the forage. Indeed, in comparison to leaves (0.17 – 0.35%) and stems (0.19 – 0.31%) [44], the content of L-DOPA is approximately 23 – 32 x higher in mucuna seeds (3.9 – 10.6%) [53], being present largely (90%) in cotyledons than the seed coat [44]. L-DOPA is potentially toxic and anti-nutritional if large amounts are ingested [86]. Whilst ruminants are generally unaffected by L-DOPA toxicity partly due to ruminal microbes being able to degrade 53% of dietary L-DOPA [56], it is possible that some ruminants are less able to degrade it, similarly to the variation in the ability to totally degrade 3,4-dihydroxy pyridine (3,4-DHP) – the ruminal metabolite of mimosine, a toxic amino acid present in the leguminous shrub *Leucaena leucocephala* – in ruminants of the same species [87]. Nonetheless, showing no undesirable effects, our results suggest that both MF and MSM can indeed be used as alternative protein sources for indigenous goats in replacement of BL.

Furthermore, our results showed that dietary MF incorporation particularly at the 100% level decreased FI whilst 100% MSM increased this parameter in goats. A possible explanation for the remarkable decrease in FI at 100% level of MF inclusion may relate to presumably high content of condensed tannins in MF. In this regard, Chikagwa-Malunga *et al*. [66] found tannins in mucuna to be concentrated in leaves and stems, with their total concentration quadratically increasing with maturity. There is a possibility that young animals used in the current study had not yet acquired the ability to utilize diets containing high levels of condensed tannins. Compared to other ruminants, goats have superior capacity to adapt to diets rich in condensed tannins [88] but this capacity is likely to be acquired with increased exposure to such diets over time [60]. However, even goats under free-range conditions will reduce intake of tannin-rich feed in order to regulate the amount of tannins in their digestive systems. In this regard, they regulate their foraging behaviour by changing the number and lengths of browsing and grazing bouts in ways that minimise condensed tannin consumption [89].

On the other hand, a possible explanation for the remarkable increase in FI at 100% level of dietary MSM inclusion may relate to the enhanced palatability induced by presumably high levels of L-DOPA in this diet. Whilst this diet had rather low fibre content, which would be expected to have increased FI, the presumably high content of the toxic L-DOPA in this diet would also be expected to have significantly decreased this parameter. However, this was not the case. Instead, the observation supports a previous study that reported high palatability of mucuna seed [90], probably induced by the presence of L-DOPA. This argument is in line with previous findings of increased DM intake in doelings fed MSM [60] and in growing sheep fed a basal diet supplemented with graded levels of mucuna pods [91]. Evidently, goats have immense liking for an L-DOPA abundant diet. Indeed, ruminants generally seem to like and consume feed containing high levels of L-DOPA without problems, though the mechanisms that enable this consumption are not understood [41].

Our study furthermore presented evidence of a diet-induced increase in hot carcass weight and dressing percentage, particularly at the 100% MSM level. This is the first study to demonstrate positive effects of dietary mucuna incorporation on carcass characteristics in indigenous goats. Nonetheless, the obtained carcass traits are within the range of literature values for native goats of similar ages [10, 17, 92]. It is only the chilling loss values of the 100% MF and 100% MSM treatment groups that stretched beyond the normal range for indigenous goats. Goat carcasses are usually estimated to have a chilling loss of 3% [93, 94]. However, Webb [17] reported that moisture and drip losses from goat carcasses are sometimes quite high (up to 8%), which may result in lesser dressing percentage values. Higher values of chilling loss (shrinkage) suggest that goats in the 100% MF and 100% MSM treatments had higher carcass weights than those in other treatments [95].

The mechanism underlying the positive effect of 100% MSM on goat carcass yield and dressing percentage might involve high rumen un-degradable protein in MSM and microbial protein synthesis [59], which supply ∼67% of the amino acids absorbed by ruminants [96]. In this connection, it is assumed that the 100% MSM dietary treatment would have contained lower levels of condensed tannins relative to the 100% MF treatment. Low (4 – 6%) dietary levels of condensed tannins increase intestinal amino acid absorption (methionine and cysteine) [97]. Also, by increasing serum glucose levels (Table 5), the 100% MSM diet would have enhanced the synthesis of glucogenic amino acids. With its assumed low tannin content [66], the 100% MSM diet would have protected these amino acids from ruminal microbial degradation [98]. Indeed, an increase in glucose concentration may be due to more by-pass protein and increased availability of glucogenic amino acids for glucose synthesis [99]. Increased post-ruminal flow of glucogenic amino acids and lipogenic moieties results in improved muscle mass accretion [100]. Also, the increased serum glucose level at 100% MSM dietary inclusion is suggestive of a MSM-induced modulation of the pattern of volatile fatty acids (VFAs) of ruminal fermentation towards greater propionate to provide energy for growing muscle [101, 102]. There is thus a need for further investigation of the ruminal VFA profile of mucuna-fed animals in future studies.

Notwithstanding, it would seem that dietary inclusion of a very high (100%) level of MF nullified the otherwise beneficial effects of condensed tannins on goat muscle accretion and carcass yield. This is probably due to very high tannin content in the 100% MF diet as evidenced by a remarkable decrease in the FI of goats fed this dietary treatment. In this regard, it is known that whilst low (4 – 6%) dietary levels of condensed tannins increase intestinal amino acid absorption [97], high (> 8%) dietary tannin concentrations can decrease animal performance due to low FI, digestibility and nitrogen absorption resulting from the destructive action of tannins on the intestinal villi and their functions [103]. As further evidence of detrimental effects of tannins in this study, almost all the carcass traits were also noticeably rather low in 100% MF-supplemented goats. This is not unexpected as botanically tannins are preferentially concentrated in mucuna leaves and stems, in comparison to pods, with their concentration being highest in mature MF [66]. The detrimental effect of high tannin diets on carcass weight and yield was previously observed in lambs fed diets supplemented with 20% and 56% *Ceratonia siliqua* (carob) pulp [104, 105]. The mechanism underlying this detrimental effect of tannins on carcass yield might entail tannin reduction of protein and amino acid availability for absorption by the animal [106].

Lastly, our results indicated no detrimental effects of dietary mucuna incorporation on the metabolic, health and immune status of goats. Instead, they showed a positive effect of mucuna, particularly 100% MSM, on serum glucose, probably caused by increased ruminal propionate production [101, 102] or by L-DOPA [107]. This contrasts a previous study by Romero *et al*. [78] who found no significant effect of 10% dietary MSM supplementation on plasma glucose concentrations in goats. Our serum glucose values were beyond the normal range of 2.78 – 4.16 mmol/L reported for healthy goats [108] and above those observed in South African indigenous Pedi goats [27]. Glucose levels lower than normal ranges are an indication of hypoglycemia while higher levels are an indication of hyperglycemia [109]. Also, our serum total protein, albumin, urea and globulin values were similar to those of indigenous goats reported by Olafadehan [110]. Further, our erythrocyte values were about 10x lower than, whereas the leucocyte values were similar to, those of Olafadehan [110]. Despite the low erythrocyte values, our goats nonetheless did not exhibit any symptoms of haemolytic anaemia. This is the first study to report effects of dietary mucuna incorporation on biochemical and haematological indices in indigenous goats. Notwithstanding, our results ascertain the safety of MF and MSM in goats. Despite the supposedly high L-DOPA content particularly in mucuna seeds [53], our data suggest that MSM can be safely used as a dietary protein supplement for indigenous goats at as high a level of incorporation as 100% without detrimental effects on animal metabolism, health and immunity.

## Conclusion

This study showed MF and MSM, in particular the latter, to be rich alternative protein sources for indigenous goats that can be used in replacement of BL. The optimal stage for harvesting and utilization of MF is between 12 and 16 WAP. Feeding mucuna caused no detrimental effects on productive performance and health status of goats but improved goat carcass characteristics, particularly when MSM was incorporated at 100% in the diet.

## Acknowledgements

We are grateful to UNESWA Transport Department for providing transport to ferry sugarcane tops and all feed ingredients from suppliers, Dalcrue Farm for the supply of sugarcane tops, Royal Swaziland Sugar Corporation for supplying molasses, Matsapha Correctional Centre for supplying broiler litter, Feedmaster (Pty) Ltd for supplying hominy chop, vitamin-mineral premix and salt, Arrowfeeds (Pty) Ltd (Eswatini) for mixing and preparation of experimental diets, Lancet Laboratories in Mbabane for the blood biochemical and haematological analyses, UNESWA Farm abattoir staff for helping with goat slaughtering and carcass measurements, and Mr. Richard Magongo in the Nutrition Laboratory for helping with chemical analyses.

## Author Contributions

Conceptualization: DMNM.

Data curation: DMNM, NPG.

Formal analysis: DMNM, NPG, AH, IVN.

Funding acquisition: DMNM.

Investigation: DMNM, NPG.

Methodology: DMNM, NPG.

Project administration: DMNM, AMD.

Resources: DMNM, NPG, AMD, AH, IVN.

Software: DMNM.

Supervision: DMNM, AMD.

Validation: DMNM.

Visualization: DMNM.

Writing – original draft: DMNM, NPG.

Writing – review & editing: DMNM, AH, IVN.

